# Employing Deep Learning Model to Evaluate Speech Information in Vocoder Simulations of Auditory Implants

**DOI:** 10.1101/2023.05.23.541843

**Authors:** Rahul Sinha, Mahan Azadpour

## Abstract

Vocoder simulations have played a crucial role in the development of sound coding and speech processing techniques for auditory implant devices. Vocoders have been extensively used to model the effects of implant signal processing as well as individual anatomy and physiology on speech perception of implant users. Traditionally, such simulations have been conducted on human subjects, which can be time-consuming and costly. In addition, perception of vocoded speech varies significantly across individual subjects, and can be significantly affected by small amounts of familiarization or exposure to vocoded sounds. In this study, we propose a novel method that differs from traditional vocoder studies. Rather than using actual human participants, we use a speech recognition model to examine the influence of vocoder-simulated cochlear implant processing on speech perception. We used the OpenAI Whisper, a recently developed advanced open-source deep learning speech recognition model. The Whisper model’s performance was evaluated on vocoded words and sentences in both quiet and noisy conditions with respect to several vocoder parameters such as number of spectral bands, input frequency range, envelope cut-off frequency, envelope dynamic range, and number of discriminable envelope steps. Our results indicate that the Whisper model exhibited human-like robustness to vocoder simulations, with performance closely mirroring that of human subjects in response to modifications in vocoder parameters. Furthermore, this proposed method has the advantage of being far less expensive and quicker than traditional human studies, while also being free from inter-individual variability in learning abilities, cognitive factors, and attentional states. Our study demonstrates the potential of employing advanced deep learning models of speech recognition in auditory prosthesis research.

## Introduction

Recent automatic speech recognition models have achieved human-level performance and impressive robustness to various forms of noise and distortion, thanks to advances in machine learning and deep learning techniques (Radford 2022). While these models were initially designed to simply match human performance, there is now growing interest in validating them as computational models of human speech perception (Weerts 2021, Rossbach, Kollmeier et al. 2022). This has the potential to offer valuable insights to the field of speech and hearing science. For instance, high-performing speech recognition models can be used to shed light on the speech cues that contribute to intelligibility across different environments, advancing the field of acoustic phonetics. Moreover, such models may also help in developing treatments for speech and hearing disorders by simulating the experience of individuals with impaired auditory systems.

This study aimed to investigate the potential of using speech recognition models to advance research in auditory implant devices, such as cochlear implants and auditory brainstem implants. Auditory implant technologies are clinically used to restore hearing in profound deafness by directly stimulating auditory neural pathways using electrodes that are surgically placed in proximity or in contact with the neural tissue. A sound coding algorithm converts incoming acoustic signal picked up by a microphone to electric stimulation of the auditory neurons. Deaf patients who use auditory implants generally benefit from their device for hearing and speech communication. However, most patients struggle participating in oral conversations in realistic acoustic environments. It is believed that further enhancement and optimization of signal processing and sound coding strategies will play a major role in improving speech perception outcomes of implant device users (Wouters, McDermott et al. 2015).

Vocoder simulation studies conducted with normal hearing human subjects have traditionally been used to model the effects of implant signal processing on speech perception as well as individual anatomy and physiology of implanted patients. Vocoders are acoustic simulations that are created with a reasonably proximate signal processing to that of implant devices (Shannon, Zeng et al. 1995). Vocoder simulations have been extensively used in the auditory implant field to evaluate the effects of number of electrodes (Shannon, Zeng et al. 1995, Dorman, Loizou et al. 1997, Shannon, Fu et al. 2004), envelope processing (Xu, Thompson et al. 2005, Souza and Rosen 2009), frequency-to-place mappings (Fitzgerald, Prosolovich et al. 2017, Jethanamest, Azadpour et al. 2017), background noise (Rosen, Souza et al. 2013, Goupell, Draves et al. 2020), channel interactions (Bingabr, Espinoza-Varas et al. 2008, Stafford, Stafford et al. 2014), etc. Though human vocoder studies have undoubtedly provided invaluable contributions to the auditory implant field, they are sometimes difficult or impossible to perform. Studies involving human subjects are generally time-consuming and costly, both in terms of subject recruitment and testing. Additionally, perception of vocoded speech spans a wide range across normally hearing listeners, and can be significantly affected by small amounts of familiarization or exposure to vocoded speech sounds (Davis, Johnsrude et al. 2005, Goupell, Draves et al. 2020). The results of human vocoder studies can be biased by individual learning abilities and cognitive factors, as well as the order of stimulus presentation and listener’s attentional state or fatigue during a testing session. These limitations can prevent researchers from being able to evaluate many different parameters with the desired precision.

Instead of testing human subjects, we employed a publicly available robust speech recognition model to evaluate the effects of vocoder parameters on speech recognition. Whisper, the open-source deep learning speech recognition model from OpenAI (Radford 2022), was used to evaluate speech information provided by vocoder simulations. We assessed the ability of the Whisper model to recognize vocoded speech signals that were generated through the simulation of various psychophysical and signal processing degradations commonly experienced by auditory implant patients. Whisper model was chosen for this study because of its advanced complex architecture and remarkable speech recognition performance in noise and different types of distortions (Radford 2022). Whisper uses the transformer language processing model with over 1.5 billion parameters (Vaswani A. 2017) and has been trained on hundreds of thousands of hours of natural speech.

The Whisper results showed a very close correspondence to the results of analogous human vocoder studies reported in the literature. The pattern of Whisper performance as a function of vocoder parameters mirrored that of similar vocoded studies with human subjects. These results support the potential of employing automatic speech recognition models for conducting vocoder studies, as a replacement for human participants. In fact, it would be almost impossible to conduct a human subject experiment with the scale of stimuli tested in this study. We evaluated Whisper’s speech recognition on roughly 270,000 sentences and 900,000 words for more than 900 vocoder and testing conditions. Evaluating this number of stimuli with human subjects would require more than 2000 hours of testing per participant, assuming that 500 sentences or words can be tested in an hour. Instead of thousands of hours of testing time spread over several months and years, the Whisper model could provide results within few hours running on the high-performance computing platforms that are currently available. These results highly support the potential of publicly available advanced speech recognition models to investigate large scale sets of parameters to evaluate or optimize signal processing for assistive hearing devices.

## Methods

### 1. Speech Recognition Model

The OpenAI Whisper (Radford 2022) is a robust multi-lingual speech recognition model, which is publicly available and open-source (https://github.com/openai/whisper/). The Whisper model is constructed based on the sophisticated transformer architecture (Vaswani A. 2017). The transformer’s ability to process context information within and across sentences has become a widely accepted and validated technique for natural language processing. Whisper has been trained on a very large speech corpus of around 680,000 hours of multilingual training data. The training set consisted of naturally produced speech tokens of various qualities and in different realistic and noisy environments. With default settings, the Whisper model assigns a probability to the possible candidates for each word in a sentence and randomly chooses among the candidates that were assigned the highest probability. This probabilistic decoding process provides flexibility in model’s decision that can result in better performance (Radford 2022). The probabilistic decoding may also result in variability in model’s performance between different runs of the model for the same speech token, which is somewhat analogues to perceptual variability and uncertainty in human speech perception.

### 2. Vocoder Processing

Noise vocoded stimuli were generated using the vocoder signal processing shown in Figure 1. The input speech signal was passed through N non-overlapping band-pass filters, and the band-passed signals were half-wave rectified and low-pass filtered to obtain envelope from each of the N channels. In the conventional vocoder processing, the resulting channel envelopes modulated narrow-band random noise signals that were filtered with the same band-pass frequency range of the corresponding channel. In some of the vocoder simulations of this study, the extracted envelopes were subjected to additional degradations before multiplying by random noise, including dynamic range reduction and quantization. Finally, the modulated noise bands were band-pass filtered by the corresponding N non-overlapping filters, and were summed to produce the final vocoder output. Band-pass and low-pass filters were implemented using sixth and fourth order Butterworth filters, respectively.

**Figure 1.**
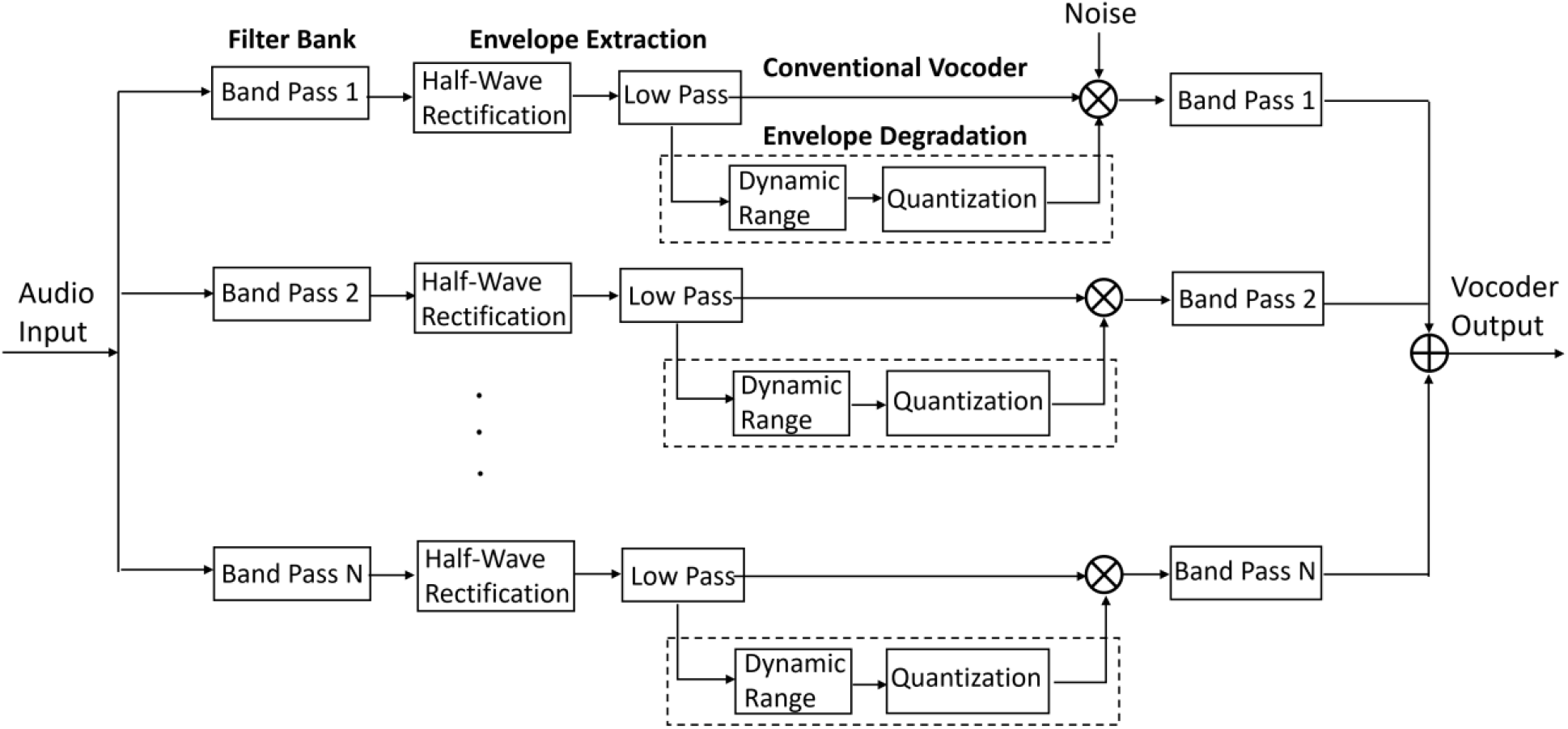
Block diagram of vocoder signal processing. The additional envelope degradation stages (dynamic range and quantization) represent manipulations in this study and not the traditional vocoder.

The vocoder processing parameters that were manipulated in this study and the rage of parameter values are shown in Table 1. These parameters intended to simulate some important signal processing and perceptual aspects of sound processing with auditory implants. The rationale for choosing each parameter is described below.

**Table I.** Vocoder parameters used for the simulations in this study

| Parameter | Range of Values |
| --- | --- |
| Number of channels | 4, 6, 8, 10, 12, 16, 22 |
| Channel Frequencies | Equal-octave bands:<br>Low cutoff frequency: 200 - 2000 Hz<br>High cutoff frequency: 2000 - 7900 Hz<br>Cochlear Ltd. Standard:<br>Low cutoff frequency: 188 Hz<br>High cutoff frequency: 7938 Hz<br>*Default: Octave bands (200-7900 Hz)* |
| Envelope Cutoff | 5 - 400 Hz<br>*Default: 100 Hz* |
| Envelope Dynamic Range | 10 -100 dB, as well as full range<br>*Default: Full range* |
| Envelope Quantized Steps | 1 – 100, as well as infinite (i.e. continuous non-quantized)<br>*Default: Infinite* |

***Number of channels*** was manipulated to simulate different numbers of active electrodes in an auditory implant device. Number of channels was varied between 4 and 22, which spans the range of active electrodes in clinical cochlear implant and auditory brainstem implant devices.

***Frequency boundary for each vocoder channel*** was chosen in two ways: 1) equal bandwidth across different channels when represented in octave, 2) frequencies mimicking the standard channel boundaries of commercial Cochlear Ltd. devices. The aim of the frequency boundary manipulations was to evaluate the effect of different approaches of determining frequency boundaries on speech recognition at different number of channels. To create equal-octave channels, the overall frequency range was converted to octaves and was divided by the number of channels. The resulted bandwidth was used to determine all the channel frequency boundaries starting from the first channel. The lowest and highest cutoff frequencies of the signal were manipulated to test the effects of the overall frequency range on the model’s speech recognition performance.

***Envelope low-pass cut-off frequency*** was manipulated to evaluate the range of envelope frequencies that provide usable speech cues. The range of tested envelope frequencies was between 5 and 400 Hz. The results have significant implications for determining the range of envelope variations that should be preserved by auditory implant devices and be accessible to device users.

***Envelope dynamic range*** was manipulated by limiting the range of envelope values in each channel. The aim of this manipulation was to evaluate the range of envelope amplitudes that contribute to speech recognition. Envelope dynamic range was manipulated by varying the low end of the envelope values, while the high end was set as the maximum envelope value across all channels for a given stimulus. Envelope values outside of the dynamic range were set to zero. The envelope dynamic range was represented as the ratio between the high and low envelope values in logarithmic scale (dB). Envelope dynamic range values tested in this study ranged between 10 and 150dB.

***The number of quantized envelope steps*** was manipulated to simulate the number of discriminable envelope steps by individual implant users. The number of discriminable steps can range from just a few to several tens among auditory implant patients (Kreft, Donaldson et al. 2004). The aim of this simulation was to evaluate the extent to which impaired sensitivity to changes in stimulus level could disrupt speech perception by implant users. Envelope quantization was implemented by rounding envelope values to the closest quantized level.

Quantized levels were obtained by dividing envelope dynamic range (in dB) into equal dB steps. Logarithmic steps were used in this study to be consistent with discriminable acoustic level steps in normally hearing human listeners. The number of quantization steps tested in this study was between 1 and 100 (Table I). It should be noted that due to the inherent nature of band-pass filtering at the final stage of vocoder processing (Figure 1), quantized envelopes would likely be slightly smeared and may not be precisely stepwise after the vocoder output was generated.

### 3. Speech testing

#### 3.1. Stimuli

We tested the Whisper model on a set of 60 AzBio Sentences and 200 CNC words. The same set of words and sentences were used for all vocoder conditions tested. Each of the 60 AzBio sentences was a separate audio file. The 200 CNC words were presented in lists of 30 words each. The reason for dividing CNC words into lists, rather than presenting one word at a time, was for models’ efficiency and accuracy. The Whisper model by default processes the audio input as 30 second clips and pads files that are shorter with zeros. Thus, the model would run much quicker when it is fed with a 30-second audio file consisting of a list of words as opposed to multiple single-word audio files. We also found that the model output was more accurate when feeding a list of words rather than playing one at a time. The vocoded sentence and word stimuli were tested in quiet and in the presence of speech-shaped noise at 5dB SNR (signal to noise ratio). Speech shaped noise was generated using the pyAcoustic python package (https://github.com/timmahrt/pyAcoustics/blob/main/pyacoustics). Since the probabilistic selection factor was used at the decoding stage of the Whisper model, the model output could vary each time it was run. We ran the Whisper model 5 times for each set of 60 sentences and 200 words at each vocoder condition, and used these repetitions to estimate a true mean and the standard error of the mean.

#### 3.2. Scoring

Since it was not feasible to grade hundreds of thousands of words and sentences by hand, we implemented automated scoring algorithms to analyze the results of sentences and words. The scoring was done by grading the output of the Whisper model, which we will refer to as *model result*, and comparing it to the correct transcription of the input audio, which we will refer to as *audio transcript*. The first step in the automatic scoring was to “clean” the model results and audio transcripts by removing all punctuations and lower-casing all letters. The model results were then compared to the corresponding audio transcripts after the cleaning phase. We implemented separate grading algorithms for sentence and word stimuli.

##### Sentence Scoring Algorithm

In this algorithm, each single sentence was graded by counting the number of common words between the model result and the audio transcript of that sentence. The total percent correct score for multiple sentences was obtained by summing the individual sentence grades and dividing by the total number of words in the set of sentences. The algorithm for grading sentences did not take the order of words into account. For example, the two sentences “Abbey skipped rocks” and “Skipped Abbey rocks” were graded as 3 correct words. This approach is consistent with how speech perception tests are commonly graded.

##### Words scoring Algorithm

The words scoring algorithm compared the model result for each 30-word list to the transcribed input of the list on a word-by-word basis. To obtain the final score, the number of words correctly recognized by the model was divided by the total number of words. One major challenge in scoring words was that the model result and the audio transcript lists didn’t always have the same number of words. The reason was that the model could miss words or interpret a single word as multiple words. We addressed this issue by using an iterative approach to finding the optimal alignment that resulted in the highest similarity between the audio transcript and the model result. The optimal word-by-word alignment between model result and audio transcript was found by iteratively sampling single words from the model result or from the audio transcript and calculating the similarity score between the remaining words.

Each sampled word was then replaced and the scoring was repeated until all the words were sampled one at a time. The sampled word associated with the highest similarity score was permanently removed and the process of sampling single words and calculating similarity score was repeated. The algorithm stopped when the number of remaining words in the model result and the audio transcript were equal. The highest score that was obtained with different alignments was used as the true correct word score.

Another issue in scoring words was to address homophones, which are words that are pronounced the same but spelled differently. An example is “whine” and “wine”. Homophones are phonetically identical and should be scored as correct when comparing the results of speech recognition model to the audio transcript. Scoring of homophones was addressed by implementing the grapheme-to-phoneme (g2p) python package (https://pypi.org/project/g2p-en/) to analyze all words that had different spellings and compared them by phonetic content rather than just spelling.

## Results

In this session, we will present the patterns of Whisper results with respect to several different vocoder parameters tested in this study. The Whisper results will be presented for words and sentences in quiet and noise. The mean and standard deviation of the 5 repetitions of each stimulus and vocoder condition are shown by symbols and error bars. The results will be compared to the analogues vocoder studies conducted with human subjects reported in the literature.

### 1. Number of channels

The Whisper model’s AzBio and CNC word scores as a function of number of channels are summarized in Figure 2. The results are shown for quiet condition as well as in the presence of speech-shaped noise at 5dB SNR. The error bars were small for many of the tested conditions, suggesting little variability in Whisper results. As expected, the performance of the Whisper model increased monotonically as the number of channels was increased from 4 to 22, and was worse in noise than in quiet. The peak performance approached ceiling in quiet, i.e. 94% for vocoded words and 99% for vocoded sentences. Scores in quiet dramatically increased from 4 channels to 6 channels, approaching a plateau at around 8-10 channels. In noise, sentence scores plateaued at around 90% at 12 channels, whereas word scores kept increasing as the number of vocoder channels was increased to 22 without reaching a clear plateau. Better recognition of sentences than single words can be explained by Whisper’s remarkable human-like ability to use contextual and linguistic information for sentence recognition. The patterns in Whisper’s performance, as function of the number of vocoded channels and the presence of noise, align with the patterns observed in vocoder studies with human subjects (Dorman, Loizou et al. 1997, Shannon, Fu et al. 2004, Grange, Culling et al. 2017).

**Figure 3.**
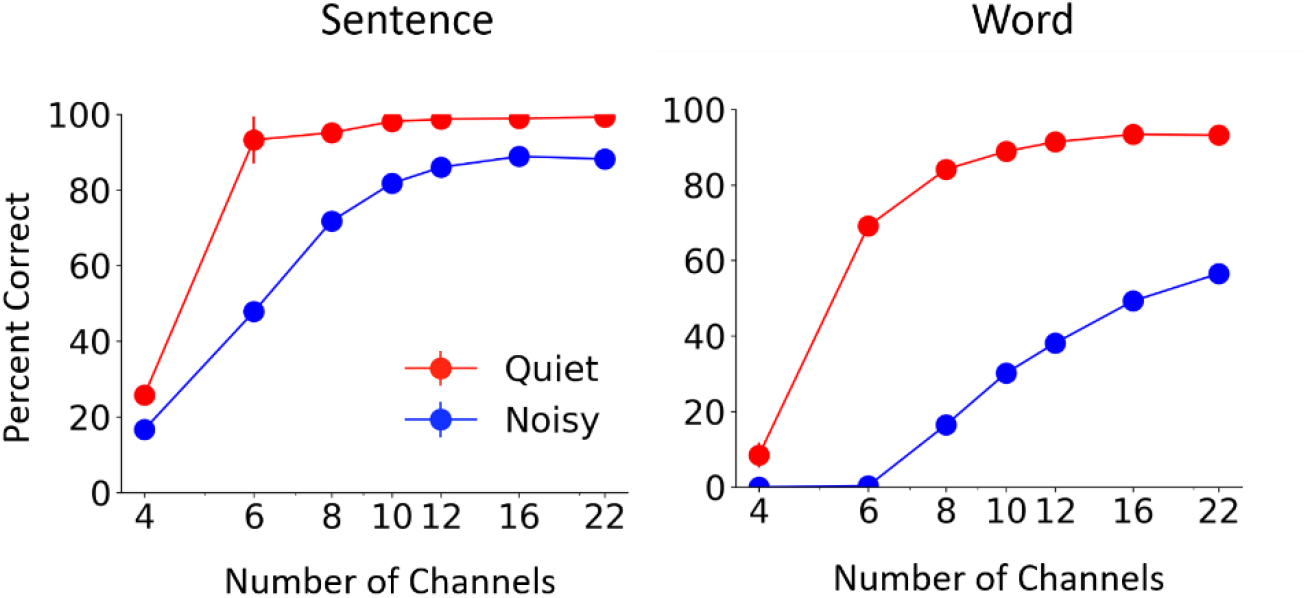
Effect of number of channels on Whisper performance in quiet and noise. Left: Mean and standard deviation of Whisper results for 5 repetitions of 60 AzBio sentences. Right: Mean and standard deviation of Whisper results for 5 repetition of 200 CNC30 words.

#### Comparison to human results

While the overall pattern of results with respect to vocoder parameters can be compared among studies, we found it challenging to compare absolute speech recognition scores between Whisper and the different human subject results reported in the literature. The reason is methodological differences between the different vocoder studies. The published human vocoder studies vary significantly in vocoder processing implementation, speech testing materials, noise type and experimental procedures. Furthermore, the duration of training or exposure to vocoder stimuli also varies between studies, with some studies providing hours of such training. All these factors could significantly affect speech perception scores when listening to vocoded speech.

To make a more valid comparison between Whisper’s performance and that of human subjects, we conducted additional model simulations under conditions similar to those of two published vocoder studies that provided details about their vocoder processing implementation and testing procedure (Grange, Culling et al. 2017, Goupell, Draves et al. 2020). Both Grange et al. and Goupell at al. used IEEE sentence material, but the type of noise was different between the two studies. The former study had speech-shaped noise while the latter used two-talker babble noise. We conducted a separate set of Whisper model simulations with IEEE sentence materials and the specific vocoder processing parameters and type of noise that were used in each of the two studies.

The results comparing Whisper to Grange at al. and Goupell et al. are shown in panels (a) and (b) of Figure 3, respectively. The data of Goupell et al. are their adult subject results obtained in the first run of testing, which is presumably minimally affected by training. The patterns of Whisper results are very similar to the results of Grange et al. and Goupell et al., in that performance improved as the number of channels and the SNR were increased. Although the patterns of whisper results agreed with human subject results, the Whisper model had slightly lower score at low number of channels (Figure 3 a) and low SNRs (Figure 3 b). Better human performance at these difficult testing conditions, at least partly, may be related to attentional, cognitive and learning abilities that are not implemented in the Whisper model. For instance, human subjects listening to vocoded speech expect to hear degraded stimuli and are prepared for the difficult task. Also, the human subjects can perceptually adapt to the degraded stimuli during the testing session. Perceptual adaptation may have particularly helped the subjects who were provided with feedback throughout the testing session (Goupell, Draves et al. 2020). Another factor that could have contributed to better human subject results in speech in noise tasks may be their awareness of the onset time of the target sentence within the background noise, enabling them to concentrate on the target speech.

**Figure 3.**
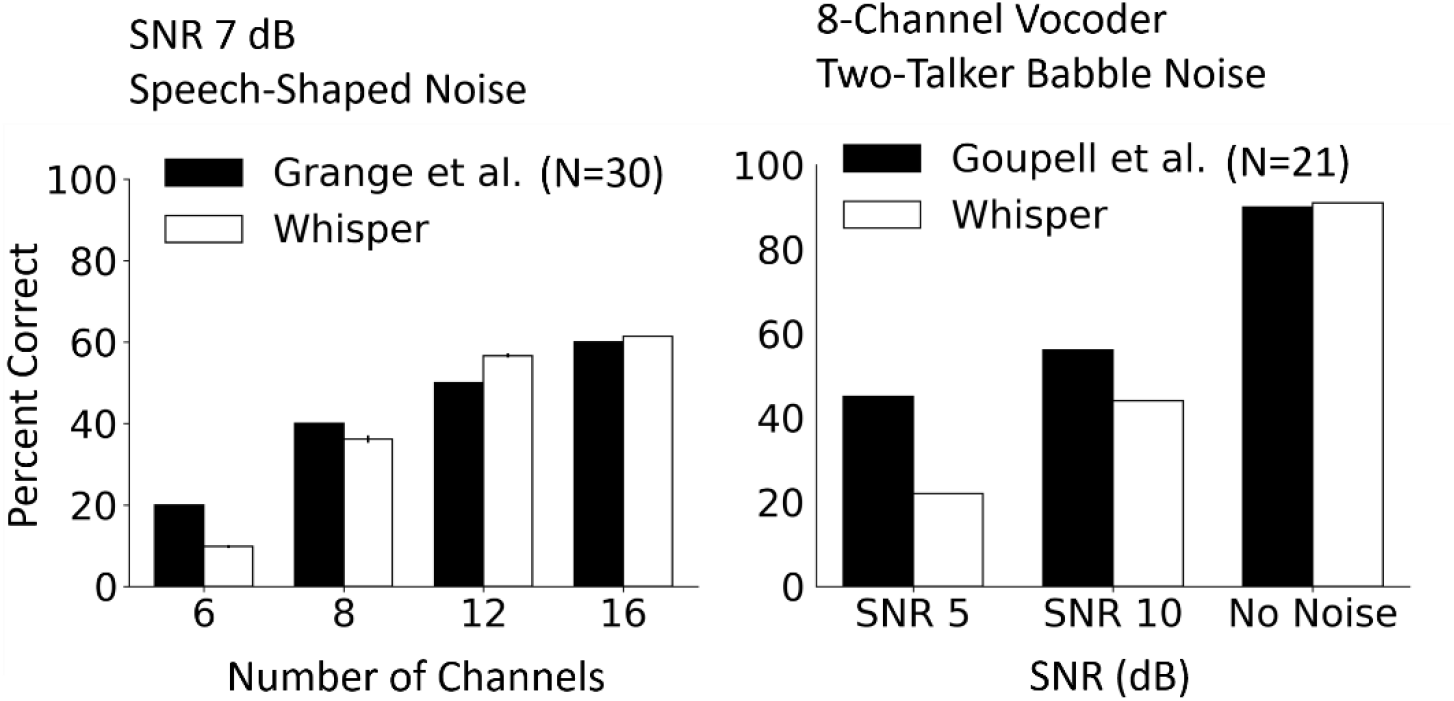
Comparison between Whisper and average human subject scores of Grange et al. (left) and Goupell et al. (right). Whisper results were obtained with similar vocoder processing parameters, sentence material, noise type and SNR to that of the reference study in each panel.

### 2. Channel frequencies

The second parameter we manipulated was the channel frequency boundaries. Frequency boundaries for different equal-octave frequency channels were manipulated by changing the overall frequency range of vocoder processing. The effects of changing the low and high cutoff of the vocoder processing are shown in Figures 4 and 5, respectively. The results showed consistent drop in performance as the staring frequency was increased (Figure 3), whereas the ending frequency didn’t have a systematic effect on the model’s performance (Figure 4). The results trends were similar for words and sentences. The detrimental effect of increasing starting frequency on speech recognition is intuitive, as higher starting frequencies discard important energy peaks (formants) in the low frequency region of the speech signal. The model’s performance dropped gradually as the starting frequency was increased up to a knee point, above which there was steeper drop in performance. The knee point was around 1000Hz for most of the vocoder conditions in quiet and was a lower frequency in noise. The drop in performance was also steeper with CNC words than with AzBio sentences.

**Figure 4.**
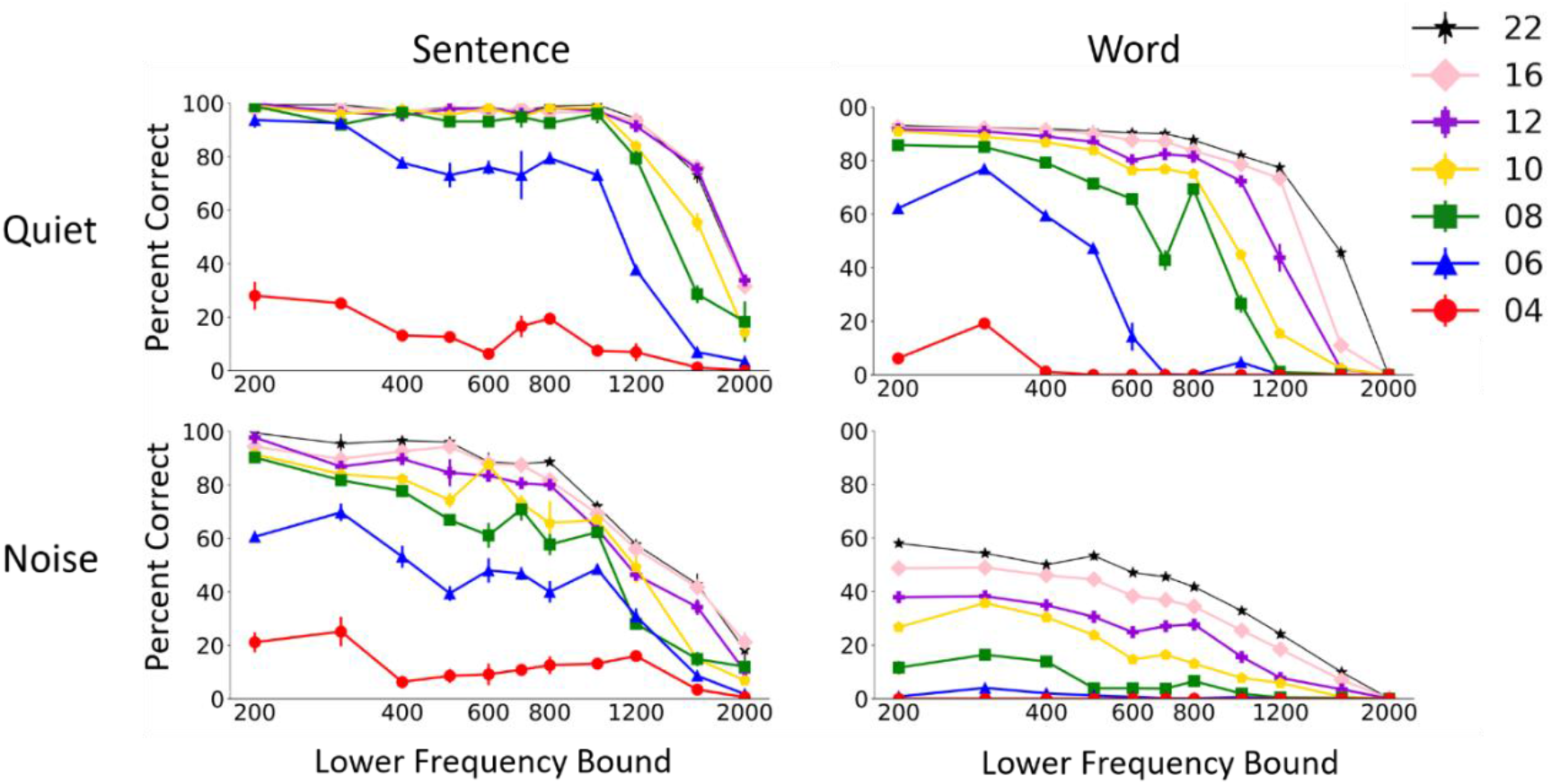
Effect of the low cutoff frequency of the signal on Whisper model performance in quiet and noise. The results show mean and standard deviation of Whisper scores for 5 repetitions of 60 AzBio sentences (left) and 200 CNC30 words (left).

**Figure 5.**
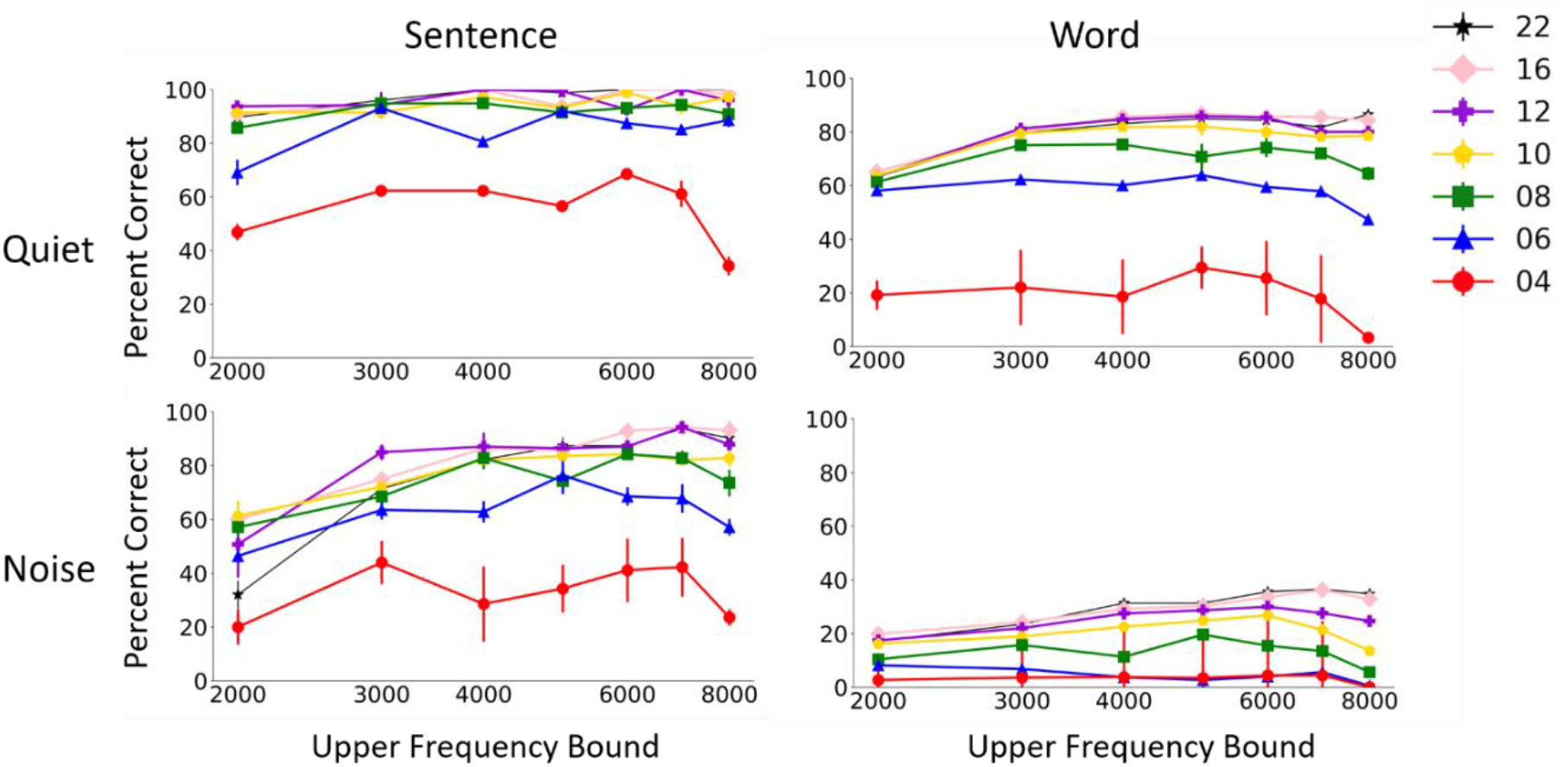
Effect of high cutoff frequency of the signal on Whisper performance in quiet and noise. The results show mean and standard deviation of Whisper scores for 5 repetitions of 60 AzBio sentences (left) and 200 CNC30 words (left).

One peculiar result is that the performance of the model showed a peak at 300Hz starting cutoff for the lower number of channels. There was also a peak in performance at ending frequency of 7000Hz. We further inspected these effects by re-testing the model with additional sets of equal-octave band frequency boundaries at starting frequencies 150, 200 and 300 Hz and ending frequencies 7000 and 7900 Hz. In addition to equal-octave bands, we also tested the standard frequency boundaries in the commercial auditory implant device by Cochlear Ltd. The results shown in Figure 6 suggest that manipulating frequency boundaries may have a large effect on recognition of noise vocoded speech, particularly for small number of channels (4, 6, and 8 channels). There was minimal effect of frequency boundary manipulation for 12 channels and above. The condition with frequency range between 300 and 7000 Hz resulted in the best model performance for all channels. The standard frequency boundaries for Cochlear outperformed most of the other tested frequency boundaries for 6 and 8 channels but showed poorer result than the other frequency conditions for 4 channels.

**Figure 6.**
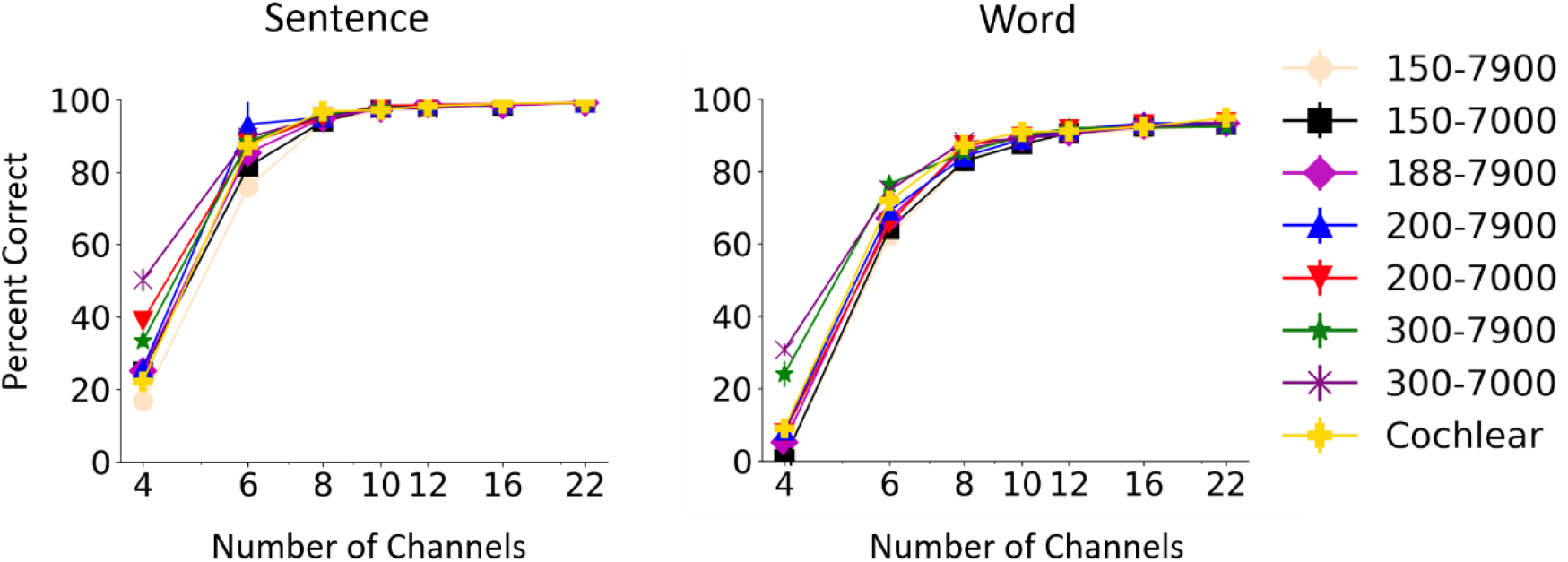
Effect of changing the frequency range of the signal on Whisper model performance at different number of channels in quiet. The results show mean and standard deviation of Whisper scores for 5 repetitions of 60 AzBio sentences (left) and 200 CNC30 words (left).

### 3. Envelope cutoff frequency

Figure 7 shows the patterns of Whisper results as a function of the envelope low-pass cutoff frequency. Whisper performance dramatically improved as envelope cutoff was increased from 5 to around 30Hz, at which point the performance plateaued regardless of the number of channels. The patterns were similar for sentence and word recognition and in quiet and noise. These results are in agreement with previous findings in human subjects, which also showed that increasing envelope cutoff beyond 30Hz had no effect on speech perception with noise vocoded speech (Souza and Rosen 2009). One interesting note is that the Whisper results in noise show a decline in performance as the low pass envelope becomes greater than 100Hz. This reduction pattern is particularly obvious for words and below 10 channels. The reason why performance in noise drops at high envelope cutoff frequencies needs further investigation but may be due to speech masking by high-frequency noise envelope that would be preserved at high cutoff frequencies.

**Figure 7.**
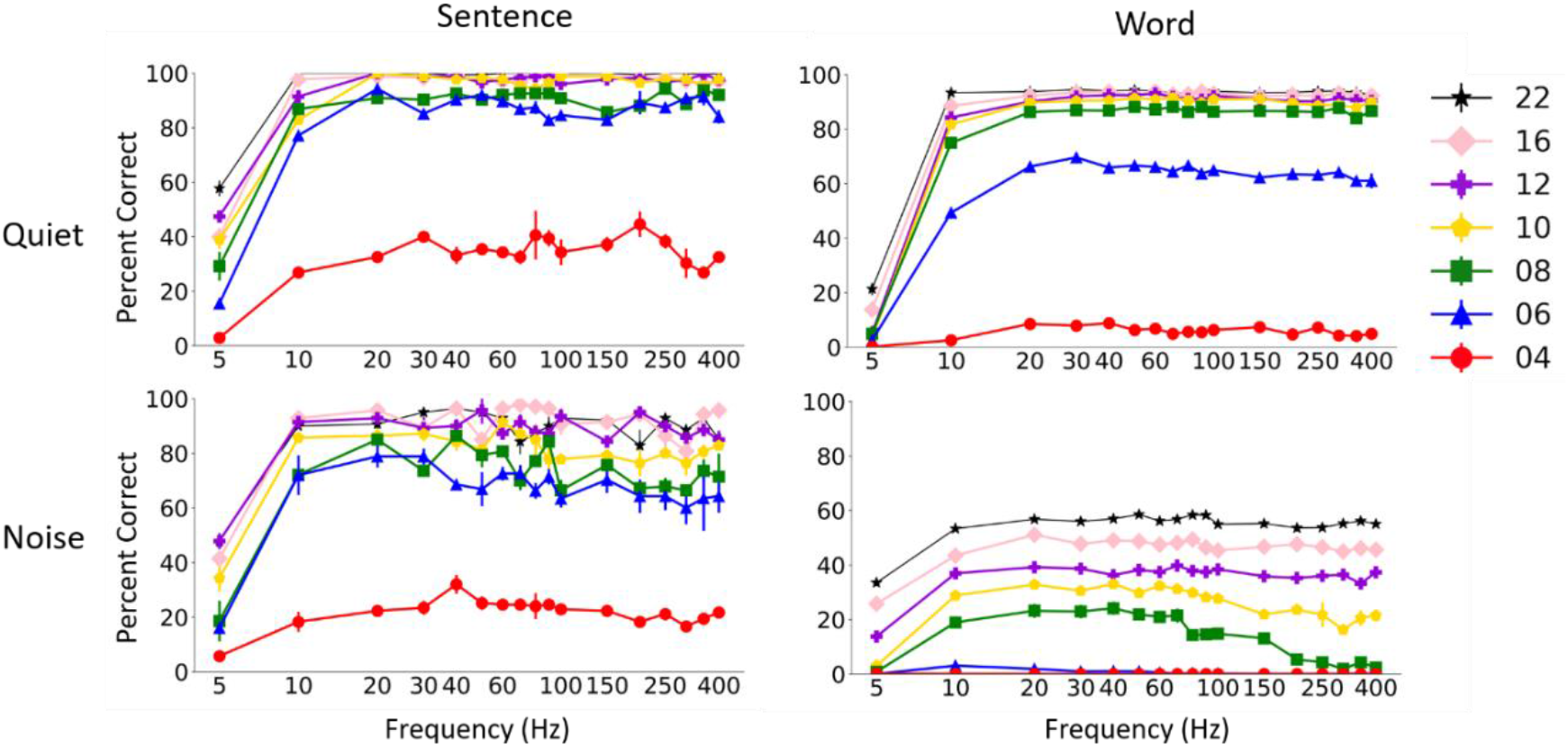
Effect of the low pass cutoff frequency of the envelope on Whisper model performance at different number of channels in quiet and noise. The results show mean and standard deviation of Whisper scores for 5 repetitions of 60 AzBio sentences (left) and 200 CNC30 words (left).

### 4. Envelope dynamic range

The results in Figure 8 suggest that envelope dynamic range could have a large effect on vocoded speech recognition. Whisper model’s sentence recognition performance considerably dropped as envelope dynamic range (the range of values below maximum) was reduced below 70 dB, particularly for words and at smaller number of channels. The patterns of changes as a function of dynamic range look similar in quiet and noise conditions. These results suggest that very low envelope levels contribute to speech recognition, particularly word recognition. To our knowledge, the effect of reducing envelope dynamic range on perception of vocoded speech by humans has not been directly evaluated. Loizou et al. showed that compressing envelope of tone vocoders after they were multiplying by output tones (at the synthesis stage of the vocoder) had a large effect on speech perception (Loizou, Dorman et al. 2000). In that study, the extracted envelope was compressed before modulating the amplitudes of tones, without removing any envelope information. This manipulation is different from our approach that reduced the range of the envelope levels. Zeng et al. evaluated the effect of input dynamic range in cochlear implant patients and found that at least 50 dB input dynamic range was needed for maximum consonant and vowel scores (Zeng, Grant et al. 2002). These results are in agreement with the results we obtained with the Whisper model in that low speech levels bear important speech cues.

**Figure 8.**
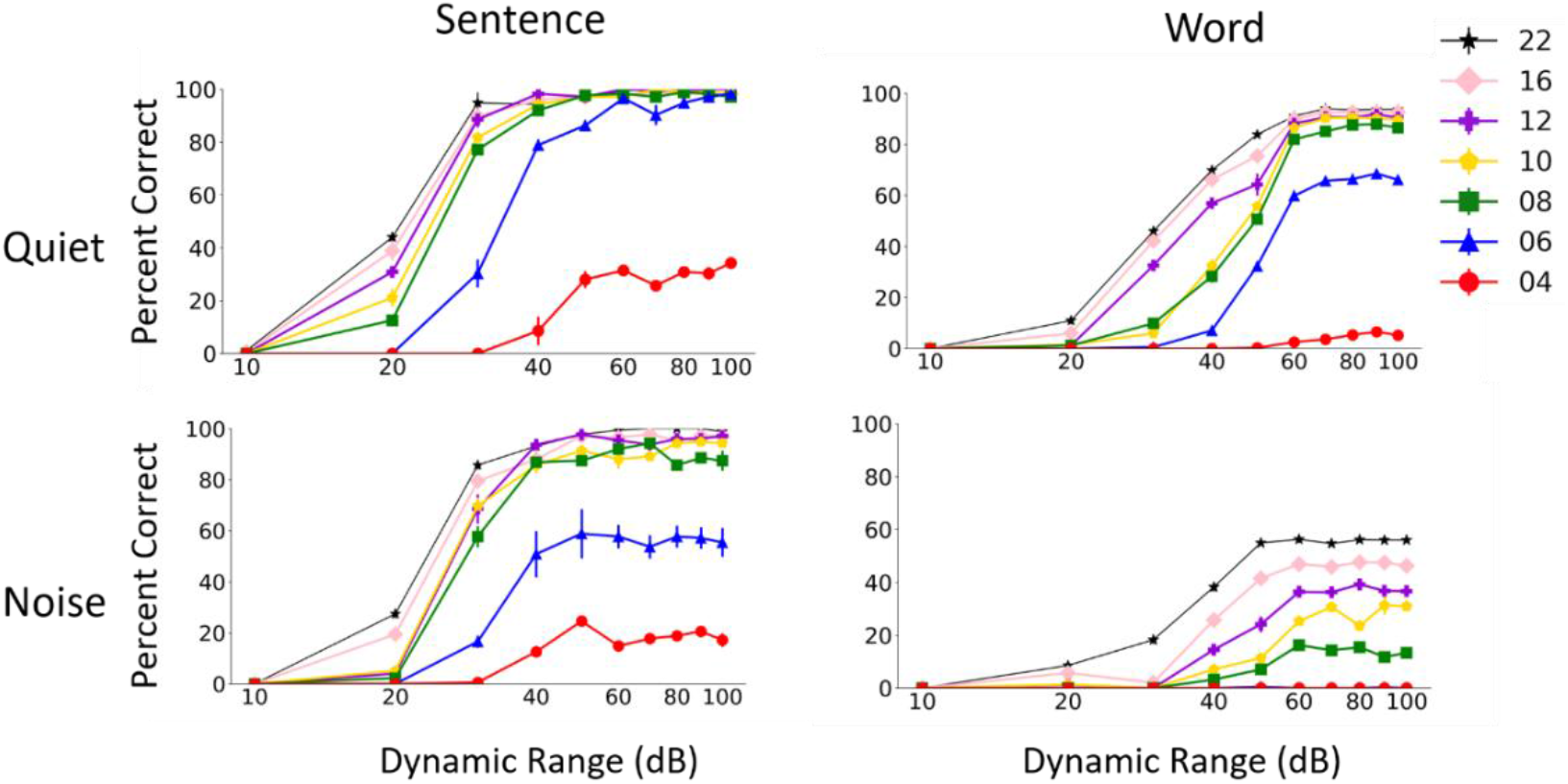
Effect of the envelope dynamic range on Whisper performance in quiet and noise. The results show mean and standard deviation of Whisper scores for 5 repetitions of 60 AzBio sentences (left) and 200 CNC30 words (left).

### 5. Envelope quantization

Figure 9 shows Whisper performance as a function of the number of quantized envelope steps. Whisper performance dramatically improved as the number of envelope steps was increased. A plateau in performance was achieved with less than 5 quantized steps for larger number of vocoder channels and in quiet. More quantized steps were needed to reach asymptotic performance in noise or with small number of channels. For most conditions, between 10 and 20 quantized steps were sufficient to achieve maximum performance. Whisper results are consistent with the results of human vocoder studies (Loizou, Dorman et al. 2000). Loizou et al. showed that human subjects listening to 16 channel vocoder required less than 5 envelope steps to reach asymptotic performance in quiet. Similar to the whisper results, the human subject results of Loizou et al. also showed that larger number of quantized steps were needed for smaller number of vocoder channels. Both Whisper and human subject results confirm that the number of discriminable steps may not be the main factor limiting speech performance of cochlear implant patients. Most cochlear implant patients have access to more than 10 discriminable steps (Nelson, Schmitz et al. 1996), which would be sufficient for near-asymptotic speech perception performance.

**Figure 9.**
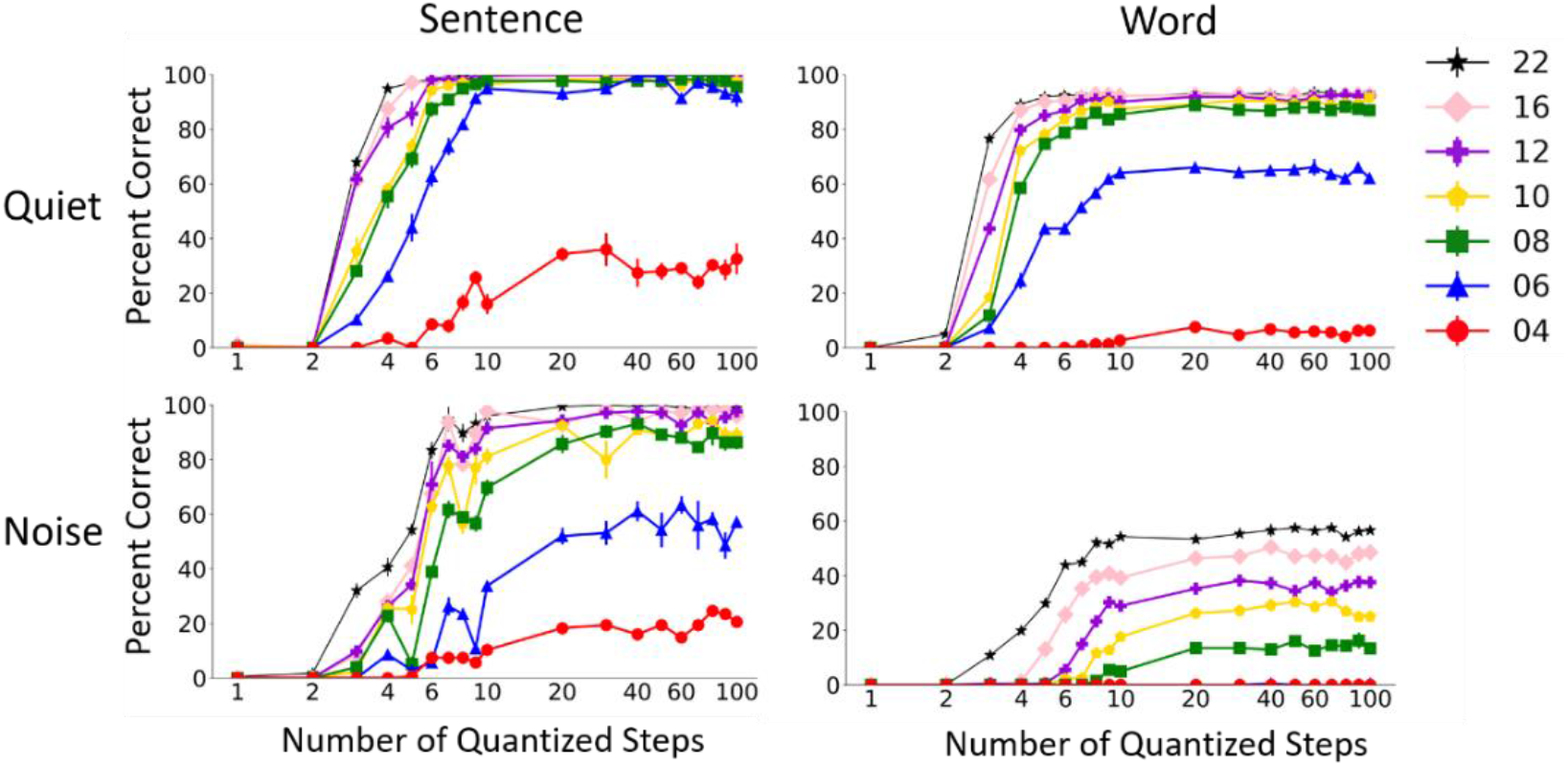
Effect of envelope quantization on Whisper performance in quiet and noise. The results show mean and standard deviation of Whisper scores for 5 repetitions of 60 AzBio sentences (left) and 200 CNC30 words (left).

## Discussion

This study employed an open-source speech recognition model released by Open AI™ to run vocoder simulations of auditory implants. Traditionally, vocoder simulations have been used to simulate implant signal processing in normal hearing listeners. The vocoder results of the model showed remarkable similarity to the published results obtained with human subjects. With only 4 channels, the model correctly recognized around 25% of AZBio sentences in quiet. The model’s sentence recognition scores dramatically increased to more than 90% at 6 channels. As expected, the model had lower CNC word scores than AzBio sentence scores. CNC word score in quiet was 10% and 60% for 4 and 6 channels, respectively. These results are generally consistent with published human subject results in quiet at different numbers of channels (Dorman, Loizou et al. 1997, Loizou, Dorman et al. 2000). It should be noted that because of methodological differences across studies, numerous factors must be taken into consideration when drawing comparisons between speech recognition findings. The factors that could significantly affect comparison between different vocoder studies are the vocoder processing parameters (e.g. channel frequency boundaries, envelope cutoff frequency, etc.), the sentence or word corpus used for testing, and the type of background noise or other environment distortions applied to the stimuli.

To provide a valid comparison between Whisper and the published human subject studies, we performed Whisper simulations using similar implementation and methodological details to two published human subject studies (Grange, Culling et al. 2017, Goupell, Draves et al. 2020). The patterns of Whisper results were very similar to the analogous human subject results, but Whisper performance was lower than human subject results in the presence of large amount of noise and for small number of channels (Figure 3). One reason that Whisper model underperforms human subjects in difficult conditions may be related to its training and deep learning architecture. Since Whisper is one of the most advanced state of the art speech recognition models to date, this possibility suggests that more technological advancements are required to achieve speech recognition models with similar phoneme and word recognition performance to humans. One other possibility that may explain better human scores is concerned with attentional and learning factors in human subject experiments. Human studies often involve various amount of training to familiarize subjects with the novel vocoded stimuli. It has been shown that even small amount of training or simply passive exposure to vocoded stimuli can significantly improve intelligibility of these sounds (Davis, Johnsrude et al. 2005, Goupell, Draves et al. 2020). In addition, the subjects in human vocoder studies are aware that they are about to hear distorted speech sounds and can benefit from that knowledge to attend to phonetic and linguistic contexts in vocoded stimuli. Human subjects’ performance might have been significantly poorer if they were completely naïve to the experimental procedure and the stimuli in each trial, which is the case for the current speech recognition models.

In this study we focused on evaluating the effects of vocoder parameters that are relevant to hearing and speech perception with auditory implants. The tested vocoder parameters directly simulate implant signal processing or perceptual aspects of the implant device users (Table I). One set of parameters we used were frequency range and frequency boundaries of vocoder channels. These parameters were used to simulate the effects of frequency-to-electrode assignment in auditory implant devices. The model results were consistent with the expectation that increasing the low cutoff frequency reduces speech perception scores. Interestingly, the results suggest that small changes in channel frequency boundaries could have a large effect on perception of vocoded speech, particularly for smaller number of channels. The next parameter we tested was envelope cutoff frequency. Envelope cutoff was manipulated to evaluate effects of signal processing as well as the individual’s ability to process rapid envelope cues. Consistent with the published human subject results, the model results showed that rapid envelope variations above 30Hz do not significantly contribute to speech understanding. Finally, we evaluated the effects of envelope dynamic range and quantized steps. These parameters were used to examine envelope amplitude cues that contribute to speech perception. The dynamic range results of the model suggested that even very low envelope levels (>60dB below maximum) can contribute to speech perception. These interesting results were rather unexpected. To the best of our knowledge, there are no published human data to directly compare to the model’s dynamic range results. The Whisper results with quantized envelope were consistent with published human studies, showing that 20 quantized steps can be sufficient for speech recognition even in noise and for low number of channels. A minimum of around 10 envelope steps was sufficient when the number of channels was greater than 8.

In general, the trend of the model results as a function of vocoder parameters agreed with the published human subject results. The vocoder results obtained with the Whisper model support the potential of employing speech recognition models in auditory implant research. These models may be used to evaluate the effects of implant signal processing parameters as well as perceptual limitations on speech perception. The focus of this paper was on using speech recognition model to evaluate vocoder simulations of auditory implants. Though vocoder simulations have played a significant role in development of auditory implant signal processing, they still cannot simulate all aspects of sound perception in an auditory implant user. For instance, the properties of the primary auditory neurons and their responses to electrical stimulation cannot be easily simulated with vocoded signals presented to acoustic hearing system. One way to leverage speech recognition models to directly study auditory implants is to modify the models to implement a biophysical model of neural activity in response to electric stimulation (Brochier, Schlittenlacher et al. 2022). Speech recognition systems typically consist of a module that emulates peripheral auditory processing in a normal ear, followed by a core neural network model of language processing. The peripheral module, originally derived from acoustic hearing, can be substituted with auditory implant physiology to directly modulate peripheral neural factors and their interactions with the implant signal. This modification to the model would allow evaluating the effects of the modeled physiological parameters of individual implant users on the model’s speech recognition performance. Other potentials of such modifications to speech recognition models could be to optimize signal processing parameters for individual implant users.

## Acknowledgments

We thank our colleagues for their valuable feedback on this work, in particular Dr. Mario Svirsky for his insightful comments on the manuscript. This study was supported by NIH/NIDCD grant R21DC020305 (PI Azadpour).

